# Improved Reference Genome for *Cyclotella Cryptica* CCMP332, a Model for Cell Wall Morphogenesis, Salinity Adaptation, and Lipid Production in Diatoms (Bacillariophyta)

**DOI:** 10.1101/2020.05.19.103069

**Authors:** Wade R. Roberts, Kala M. Downey, Elizabeth C. Ruck, Jesse C. Traller, Andrew J. Alverson

## Abstract

The diatom, *Cyclotella cryptica*, is a well-established experimental model for physiological studies and, more recently, biotechnology applications of diatoms. To further facilitate its use as a model diatom species, we report an improved reference genome assembly and annotation for *C. cryptica* strain CCMP332. We used a combination of long- and short-read sequencing to assemble a high-quality and contaminant-free genome. The genome is 171 Mb in size and consists of 662 scaffolds with a scaffold N50 of 494 kb. This represents a 176-fold decrease in scaffold number and 41-fold increase in scaffold N50 compared to the previous assembly. The genome contains 21,250 predicted genes, 75% of which were assigned putative functions. Repetitive DNA comprises 59% of the genome, and an improved classification of repetitive elements indicated that a historically steady accumulation of transposable elements has contributed to the relatively large size of the *C. cryptica* genome. The high-quality *C. cryptica* genome will serve as a valuable reference for ecological, genetic, and biotechnology studies of diatoms.

**Data available from:** NCBI BioProjects PRJNA628076 and PRJNA589195

## Introduction

The diatom *Cyclotella cryptica* Reimann, J.C.Lewin & Guillard has a range of properties that have made it a valuable experimental model in studies dating back to the 1960s (Lewin and Lewin 1960). *Cyclotella cryptica* can grow across a broad range of salinities, and its responses to altered salinity offer opportunities to study several important aspects of diatom biology. For example, salinity shifts can induce gamete production (Schultz and Trainor 1970) and cause cells to alternate between cell wall morphologies resembling *C. cryptica* and the closely related freshwater species, *Cyclotella meneghiniana* Kützing (Schultz 1971). Later studies demonstrated the utility of *C. cryptica* for understanding cell wall morphogenesis in diatoms (Tesson and Hildebrand 2010). *Cyclotella cryptica* has other properties that make it an attractive candidate for biotechnology applications, including the ability to grow heterotrophically (Hellebust 1971; White 1974; Pahl *et al.* 2010) and produce high levels of lipids for use as biofuels or nutraceuticals (Roessler 1988; Traller and Hildebrand 2013; Slocombe *et al.* 2015).

A draft genome assembly for *C. cryptica* revealed a large, gene- and repeat-rich genome (Traller *et al.* 2016). The genome was sequenced without the benefit of long-read sequencing platforms, which enable short contigs—particularly those containing repetitive DNA—to be joined into large contiguous scaffolds. Consequently, the version 1.0 genome assembly of *C. cryptica* was highly fragmented, with most fragments measuring <1 kb in length. Although the gene space appeared to be well characterized and the size accurately estimated, highly fragmented assemblies can suffer from overestimation of gene number (Denton *et al.* 2014) and hinder insights into genome structure. It is also challenging to fully characterize intergenic regions, which hold noncoding RNAs, promoter regions, and allow comparisons of genomic synteny across species. This is especially challenging for historically understudied groups, such as diatoms, in which the pace of genomic sequencing has lagged behind other groups such as animals and flowering plants. The relatively small number of sequenced genomes from distantly related diatom species gives the impression each newly sequenced diatom genome contains a large fraction of unique, species-specific sequence. As small fragments, the origin and identity of these sequence fragments are especially challenging to characterize. Diatoms maintain intimate relationships with bacteria both in nature (Amin *et al.* 2012) and in cell culture (Johansson *et al.* 2019). In addition, some diatoms may have incorporated bacterial genes into their genomes (Bowler *et al.* 2008). With long contiguous scaffolds, the proximal source of bacterial-like genes should be much easier to determine in diatom genome assemblies that contain a mix of DNA from both the diatom and its associated bacteria.

We combined short and long sequencing reads to produce a more contiguous version 2.0 genome assembly for *C. cryptica* CCMP332. The addition of long direct sequencing reads allowed us to improve the gene models, remove contaminant sequences, and better characterize the structure of this relatively large genome. As a result, the improved assembly provides a better resource for functional studies of *C. cryptica* such as genome-enabled reverse genetics and readmapping for resequencing and experimental transcriptomics.

## Materials And Methods

### Strain information and sequencing

We acquired *Cyclotella cryptica* strain CCMP332 from the National Center for Marine Algae and Microbiota (NCMA). This strain was originally isolated from Martha’s Vineyard, MA, USA, by R. Guillard in 1956. We grew the culture in L1 marine medium (Guillard 1975) at 22°C on a 12:12 light: dark cycle.

We harvested non-axenic cells during late exponential-phase growth, filtered them through 5.0 μm Millipore membrane filters to reduce the bacterial load, rinsed the cells from the filter before pelleting them by centrifugation at 2500 x g for 10 minutes, and stored the cell pellets at −80°C. We extracted DNA using the DNeasy Plant Kit (Qiagen) or a modified CTAB protocol (Doyle and Doyle 1987). For the CTAB protocol, we resuspended cell pellets in 3× CTAB buffer (CTAB, 3% w/v; 1.4 M NaCl; 20 mM EDTA, pH 8.0; 100 mM Tris-HCl, pH 8.0; 0.2% β-mercaptoethanol), disrupted them by vortexing briefly with 1.0 mm glass beads, and incubated them at 65°C for 1 hour. We then extracted the DNA twice with 1 × volume of 24:1 chloroform:isoamyl alcohol and precipitated the DNA with 1× volume of isopropanol and 0.8× volume of 7.5 M ammonium acetate. We assessed the quality and quantity of the DNA with 0.8% agarose gels, a Nanodrop 2000 (Thermo Fisher Scientific), and a Qubit 2.0 Fluorometer (dsDNA BR kit; Thermo Fisher Scientific).

For DNA samples with high molecular weight and sufficient quantity (1-3 μg), we prepared libraries for long-read sequencing using the ligation sequencing kit SQK-LSK108 (Oxford Nanopore Technologies, ONT). We sequenced the libraries using the MinION platform with FLO-MIN106 (R9.4.1) flowcells (Table S1). After sequencing, we used Guppy (version 2.3.5) (ONT) with default settings to convert raw signal intensity data into base calls. We kept all nanopore reads with a length greater than 500 bp and trimmed them for adapter sequences with NanoPack (De Coster *et al.* 2018). We used Canu (version 1.7) (Koren *et al.* 2017) to correct base calls in the low quality of nanopore raw reads.

We prepared short-read Illumina sequencing libraries using the Kapa HyperPlus Kit (Roche) with 300–400 bp insert sizes and barcoded the libraries with dual indices. These libraries were sequenced using the Illumina HiSeq4000 at the University of Chicago Genomics Facility. Twelve libraries were sequenced for 50 bp single end (SE) reads and 3 libraries were sequenced for 100 bp paired-end (PE) reads (Table S1). We quality trimmed the short-reads using Trimmomatic (version 0.36) (Bolger et al. 2014) with options ‘ILLUMINACLIP:TruSeq3-PE-2.fa:2:30:10 LEADING:3 TRAILING:3 SLIDINGWINDOW:4:15 MINLEN: 50’.

### Genome assembly, error correction, and scaffolding

K-mer based analyses estimated the haploid genome size of *C. cryptica* to be 161.7 Mb (Traller *et al.* 2016). We used an estimated genome size of 165 Mb for genome assembly. We assembled the metagenome (*C. cryptica* and associated bacteria) of our culture using Flye (version 2.4.2) (Kolmogorov *et al.* 2019b, 2019a) with the raw nanopore reads and options ‘--meta --plasmids --iterations 1 --genome-size 165m’. We then mapped the corrected nanopore reads back to the assembled contigs with Minimap2 (version 2.10-r761) (Li 2018), using the settings recommended for nanopore reads. We used these mappings for error correction of the initial draft assembly with Racon (version 1.3.3) (Vaser *et al.* 2017) using default settings. We then used the r941_flip935 model in Medaka (version 0.8.1) (https://github.com/nanoporetech/medaka) for a second round of error correction using the Racon-corrected contigs and the corrected nanopore reads. After contig correction, we separately aligned the SE and PE Illumina reads to the corrected contigs using BWA-MEM (version 0.7.17-r1188) (Li and Durbin 2009). We merged and sorted the alignment BAM files using SAMTOOLS (version 1.9) (Li *et al.* 2009) and used the merged BAM file for sequence polishing of all variant types (‘--changes --fix all’) with Pilon (version 1.23) (Walker *et al.* 2014). We performed three iterative rounds of Illumina read mapping and Pilon polishing.

We scaffolded the polished contigs using the corrected nanopore sequences with SSPACE-LongRead (version 1-1) (Boetzer and Pirovano 2014), requiring three overlaps to connect contigs (‘-l 3’). We used the corrected nanopore reads to extend contigs and fill scaffold gaps using LR_Gapcloser (Xu *et al.* 2019) with default settings and a total of ten iterative rounds. Finally, we aligned the corrected nanopore reads to the scaffolds with Minimap2 and used these alignments to remove redundant scaffolds from the assembly using Purge Haplotigs (version 1.0.0) (Roach *et al.* 2018). We evaluated each stage of the assembly with QUAST (version 5.0.0) (Gurevich *et al.* 2013) and BUSCO (version 4.0.6; genome mode, eukaryote_odb10 dataset) (Simão *et al.* 2015) (Table S2).

### Contaminant identification and removal

We used the Blobtools pipeline (version 1.1.1) (Laetsch and Blaxter 2017) to identify and remove contaminant scaffolds. Blobtools uses a combination of taxonomic assignment, GC percentage, and read coverage to identify contaminants. We assigned the taxonomy of each scaffold from a Diamond BLASTX search (version 0.9.21) (Buchfink *et al.* 2015) against the UniProt Reference Proteomes database (release 2019_06) (UniProt Consortium 2018) using options ‘--max-target-seqs 1 --sensitive --evalue 1e-25 --outfmt 6’. We estimated read coverage using all reads (corrected nanopore and Illumina) mapped to the scaffolds with Minimap2, and merged and sorted the alignments using SAMTOOLS. We flagged and removed scaffolds that met the following criteria: (1) taxonomic assignment to bacteria, archaea, or viruses, (2) low GC percentage indicative of organellar scaffolds, and (3) no taxonomic assignment for scaffolds <1 kb in length. Following the removal of the contaminant sequences, we performed an additional two rounds of Pilon polishing as described above.

### RNA sequencing and assembly

We used RNA-seq reads and transcriptome assemblies for *C. cryptica* CCMP332 from Nakov et al. (2020). For that study, total RNA was extracted from cells grown in five different salinity treatments (0, 2, 12, 24, 36 parts per thousand) using the RNeasy Plant Kit (Qiagen), and 15 Illumina libraries were prepared using the Kapa mRNA HyperPrep kit and sequenced on the Illumina HiSeq2000 platform at the Beijing Genomics Institute.

### Gene annotation

We used the MAKER software package (version 2.31.10) to identify protein-coding genes in the genome (Cantarel *et al.* 2008; Holt and Yandell 2011). We used the *C. cryptica* transcriptome as expressed sequence tag (EST) evidence (est2genome=1) and the protein sequences from *Cyclotella nana, Thalassiosira oceanica, Phaeodactylum tricornutum*, and *Fragilariopsis cylindrus* as protein evidence (protein2genome=1) for the MAKER pipeline. Protein sequences were downloaded from the Joint Genome Institutes (JGI) PhycoCosm resource (https://phycocosm.jgi.doe.gov/phycocosm/home; last accessed 2 Jan 2020). We also allowed MAKER to predict single exon genes (single_exon=1) and search for alternative splicing (alt_splice=1). Repetitive elements identified during the repeat analysis (see below) were used to mask the repetitive regions for this analysis. After the first round of MAKER using EST and protein evidence, we used the predicted genes with annotation edit distance (AED) scores less than 0.5 to train gene prediction models in SNAP (version 2006-07-28) (Korf 2004) and Augustus (version 3.3.2) (Stanke *et al.* 2008). We then performed two subsequent rounds of MAKER annotation using the trained SNAP and Augustus models. We retrained SNAP after the second round of MAKER. We evaluated the completeness and quality of the MAKER proteins after each round using BUSCO (protein mode against the eukaryota_odb9 dataset) and AED scores (Table S3).

To identify protein families, domains, and gene ontology (GO) terms, we searched the predicted protein sequences against the Pfam (version 32.0) (El-Gebali *et al.* 2019), PRINTS (version 42.0) (Attwood *et al.* 2012), PANTHER (version 14.1) (Thomas *et al.* 2003), SMART (version 7.1) (Letunic *et al.* 2012), SignalP (version 4.1) (Petersen *et al.* 2011), and TMHMM (version 2.0c) (Krogh *et al.* 2001) databases using InterProScan (version 5.36-75.0) (Jones *et al.* 2014). We also searched the proteins against the SwissProt (release 2019_06) and UniProt Reference Proteomes (release 2019_06) databases using NCBI BLASTP (version 2.4.0+) (Camacho *et al.* 2009) using options ‘-evalue 1e-6 -outfmt 6 -num_alignments 1 -seg yes - soft_masking true -lcase_masking -max_hsps 1’.

We predicted non-coding RNAs (ncRNAs) in the genome using Infernal (version 1.1.2) (Nawrocki and Eddy 2013) against the Rfam database (version 14.1) (Kalvari *et al.* 2018). We used tRNAscan-SE (version 2.0.5) (Chan and Lowe 2019) for tRNA annotation and RNAmmer (version 1.2) (Lagesen *et al.* 2007) for rRNA annotation.

Chloroplast and mitochondrial genomes were annotated with GeSeq (Tillich *et al.* 2017). Gene and inverted repeat boundaries from GeSeq were manually curated as necessary by comparison to annotations from sequenced diatom organellar genomes.

### Repetitive element annotation

We built custom repeat libraries to identify repetitive elements across the genome. We searched for long terminal repeat (LTRs) retrotransposons using the program LTRharvest (version 1.5.8) (Ellinghaus *et al.* 2008) with options ‘-minlenltr 100 -maxlenltr 6000 -mindistltr 1500 -maxdistltr 25000 -motif tgca -similar 85 -mintsd 5 -maxtsd 5 -vic 10’. We filtered the candidate LTRs from LTRharvest using LTRdigest (version 1.5.8) (Steinbiss *et al.* 2009) to keep elements with polypurine tracts (PPT) and primer binding sites (PBS) inside the predicted LTR sequence region. We further filtered LTR elements using Perl scripts (Campbell *et al.* 2014) to remove elements with nested insertions and select representative (exemplar) elements. We identified miniature inverted transposable elements (MITEs) with MITE-Hunter (Han and Wessler 2010). We then masked the genome with the combined LTR and MITE libraries using RepeatMasker (version 4.0.5) (https://www.repeatmasker.org). After masking, we identified other repetitive elements using RECON (version 1.08) (Bao and Eddy 2002) and RepeatScout (version 1.06) (Price *et al.* 2005) as implemented within the RepeatModeler package (version 2.0) (Flynn *et al.* 2020). We combined all candidate exemplar elements and searched them against the UniProt Reference Proteomes database with NCBI BLASTX using settings ‘-evalue 1e-10 -num_descriptions 10’. We removed elements from the final repeat library that contained overlaps with any predicted proteins using ProtExcluder (version 1.2) (Campbell *et al.* 2014).

We used RepeatMasker and the final repeat library to annotate the repetitive elements in the genome. We ran RepeatMasker with the NCBI RMBLAST (version 2.6.0+) search engine (‘-e ncbi’), the sensitive option (‘-s’), and the ‘-a’ option to obtain the alignment file. We then used the provided parseRM.pl script (version 5.8.2) (downloaded from https://github.com/4ureliek/Parsing-RepeatMasker-Outputs) on the alignment files from RepeatMasker to generate the repeat landscape with the ‘-l’ option (Kapusta *et al.* 2017). This script collects the percent divergence from the repeat library for each TE element, correcting for higher mutation rates at CpG sites and using Kimura 2-Parameter distance output by RepeatMasker. The percent divergence to the repeat library is a proxy for age (older TE elements will have accumulated more nucleotide substitutions), and the script splits TEs into bins of 1% divergence.

### Assembly comparisons

We downloaded the *C. cryptica* version 1.0 genome assembly and gene models from http://genomes.mcdb.ucla.edu/Cyclotella/download.html. We downloaded the *P. tricornutum* version 2.0, *F. cylindrus* version 1.0, and *C. nana* version 3.0 genome assemblies from GenBank (accession numbers available in Table 1). To compare these genomes against the *C. cryptica* version 2.0 assembly, we performed QUAST and BUSCO analyses as described above.

We identified putative contaminant scaffolds in the *C. cryptica* version 1.0 assembly using the same Blobtools procedure described above and with read-mapping information from our Illumina reads. To compare functional information between the *C. cryptica* versions 1.0 and 2.0 annotations, we searched the proteins from the version 1.0 assembly against the Pfam, PRINTS, PANTHER, SMART, SignalP, and TMHMM databases using InterProScan. We also searched the proteins against the SwissProt and UniProt Reference Proteomes databases using NCBI BLASTP.

To assess overlap between the *C. cryptica* versions 1.0 and 2.0 annotations, we aligned predicted protein sequences from the two genomes to one another with NCBI BLASTP with options ‘-evalue 1e-6 -max_target_seqs 1 -max_hsps 1 -outfmt 6’. We parsed these results to count those with the same length (qlen = slen), those with 100% identity (pident = 100), those with high similarity (pident ≥ 90), and those with full alignment lengths (qcovs = 100).

### Data availability

The genome assembly and sequence data are available from NCBI BioProject PRJNA628076. RNAseq data are available through the NCBI Short Read Archive under BioProject PRJNA589195. A genome browser and gene annotations are available through the Comparative Genomics (CoGe) web platform (https://genomevolution.org/coge/) under genome ID 57836. File S1 contains Tables S1–S6. File S2 contains Figures S1–S3. File S3 contains the genome annotation GFF3, protein fasta, and transcript fasta files. File S4 contains the non-coding RNA annotation files. File S5 contains the repeat element annotation files.

## Results And Discussion

### Genome assembly

We sequenced five libraries on the MinION platform and base called over 5.9 million reads that totaled 9.42 Gb of sequence, where the read length N50 was 4.4 kb and the median quality score per read was 9.9 (Table S1 and Figure S1). After trimming and filtering, we used a total of 2,941,466 nanopore reads with a median quality score per read of 10.1 for genome assembly (Figure S1). We also sequenced 15 short-read libraries on the Illumina platform which provided nearly 450 million reads, totaling 35.3 Gb of sequence data (Table S1). Transcriptome sequencing of 15 libraries were assembled into 35,726 transcripts and used for genome annotation (Nakov et al. 2020).

On average, each position in the version 2.0 genome was covered by 152 reads, including both the Nanopore (74×) and Illumina (78×) reads. The new assembly represents a substantial improvement over the original version 1.0 genome assembly, which was generated from Illumina short-read sequencing data only. The application of long-read sequencing data resulted in several important changes or improvements, including: (1) an increase in the estimated genome size, from 161.7 Mb to 171.1 Mb; (2) a 176-fold decrease in the number of scaffolds, from 116,815 in version 1.0 to 662 in version 2.0; (3) a 41-fold increase in the scaffold N50, from 12 kb in version 1.0 to 494 kb in version 2.0; (4) a substantial decrease in the number of N’s linking contigs into scaffolds, from 5,360 N’s per 100 kb in version 1.0 to 52 N’s per 100 kb in version 2.0; and (5) increased detection of conserved eukaryotic orthologs, from 183/255 (72%) complete BUSCO genes in version 1.0 to 191/255 (75%) in version 2.0 (Table 1 and Figure 1). The BUSCO count for *C. cryptica* is now on par with those of the model diatoms, *C. nana* (75%), *F. cylindrus* (78%), and *P. tricornutum* (78.5%) (Table 1 and Figure 1).

The plastid genome assembly was 129,328 bp in total length with 2,707× coverage, which did not differ significantly from the 129,320 bp plastid genome size reported in the version 1.0 assembly (Table 1). The mitochondrial genome was assembled to a total size of 46,485 bp with 2,520× coverage, which was nearly 12 kb shorter than the 58,021 bp genome assembled previously (Table 1). This 12 kb difference reflects the size of the complex repeat region present in many diatom mitochondrial genomes (Oudot-Le Secq and Green 2011). We were able to fully span this region with long sequencing reads.

**Table 1.**
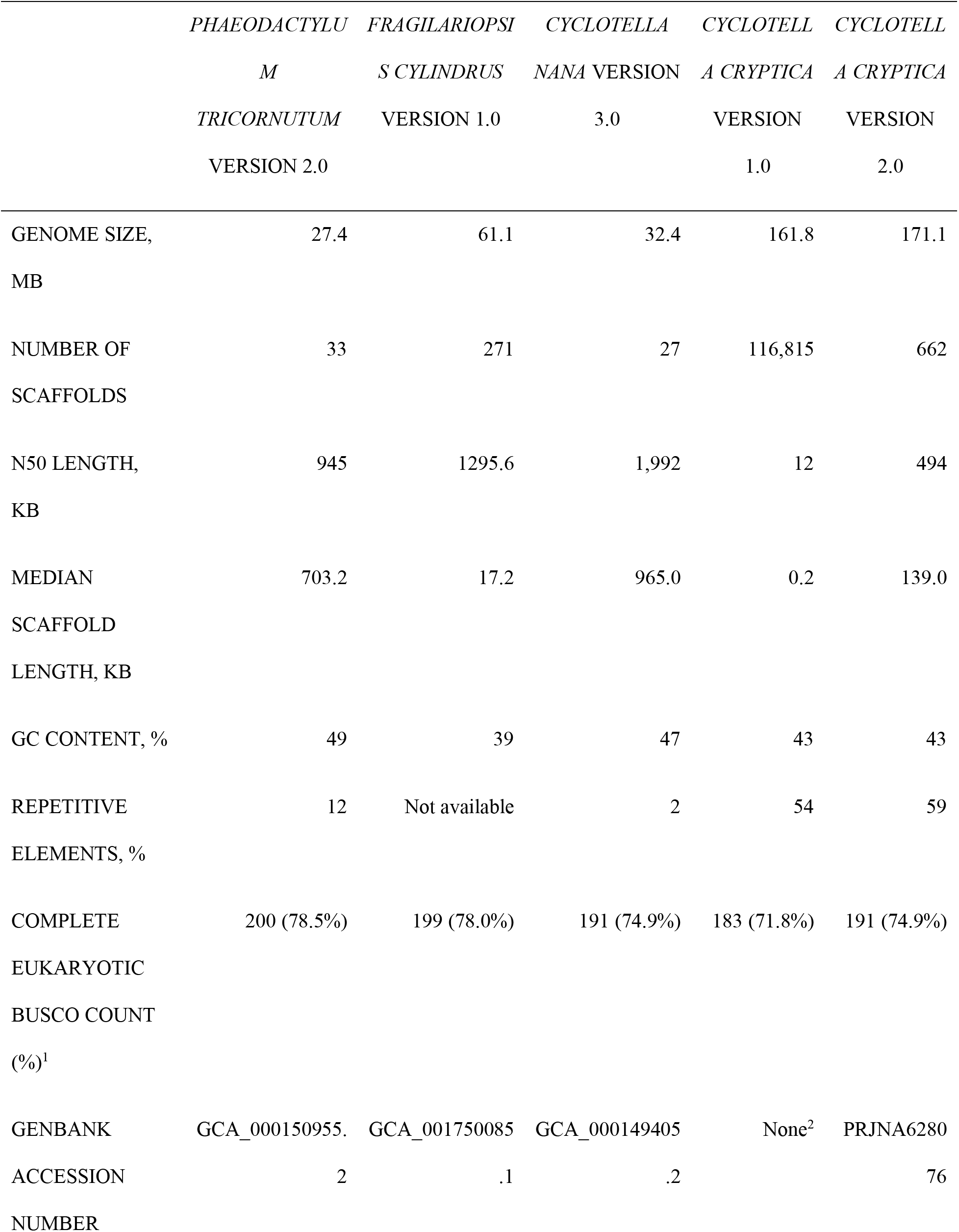
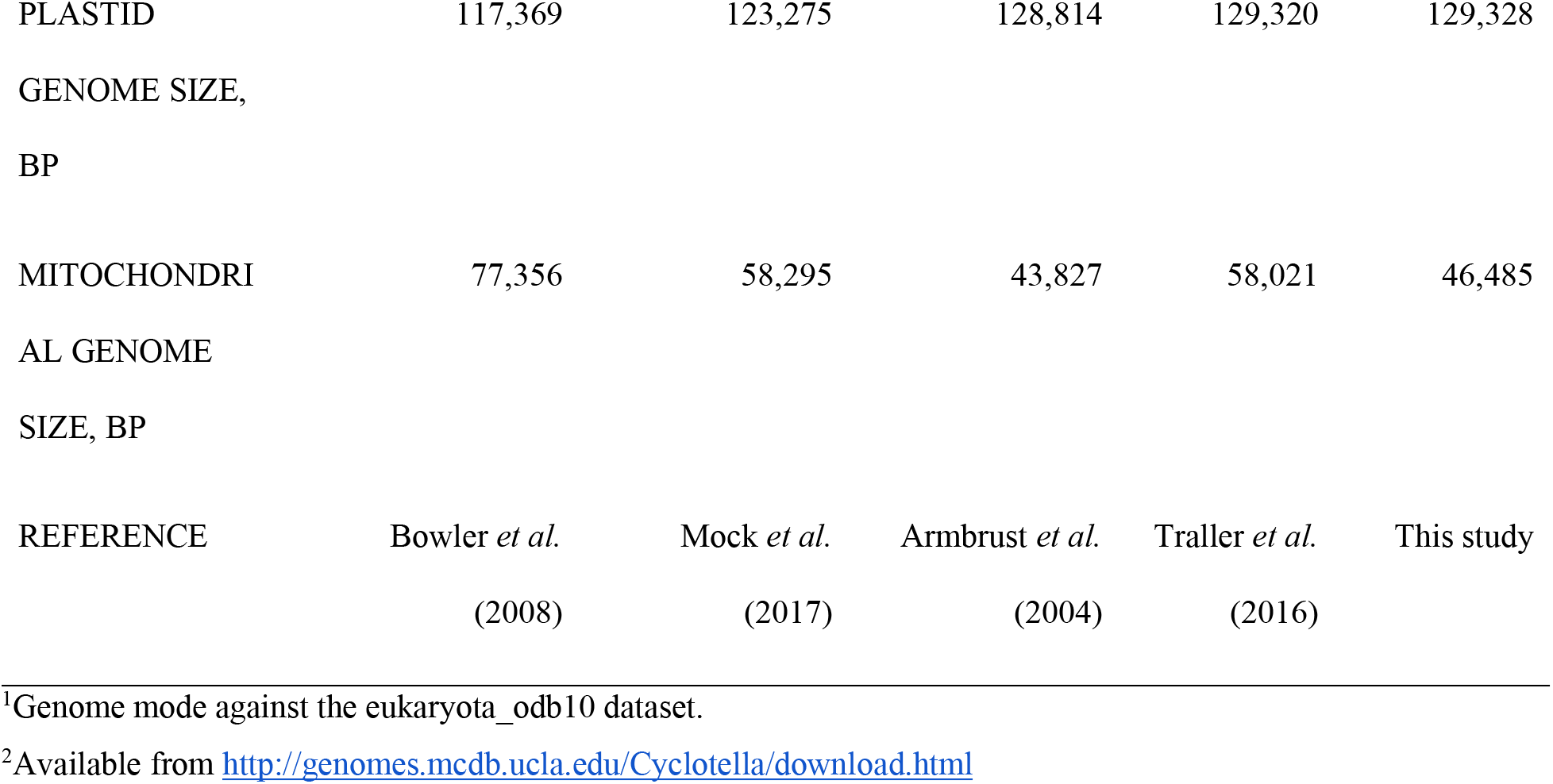
Genome characteristics for *P. tricornutum, F. cylindrus, C. nana*, and *C. cryptica*.

**Figure 1.**
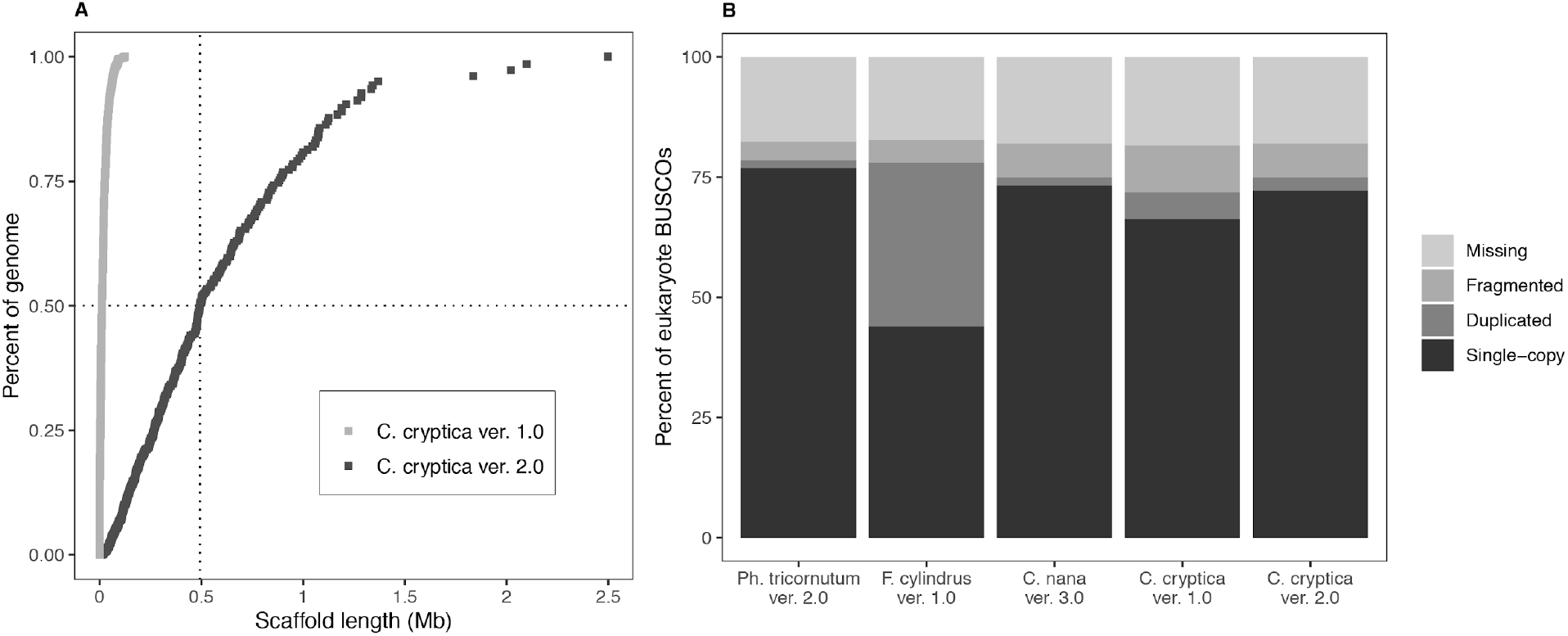
Improved genome assembly for *Cyclotella cryptica.* (A) Cumulative scaffold length and N50 comparison in the version 1.0 and version 2.0 assemblies. Summary statistics for each assembly are given in Table 1. (B) BUSCO analysis of selected diatom genomes using the set of 255 conserved eukaryotic single-copy orthologs. Bars show the proportions of genes found in each assembly as a percentage of the total gene set.

### Bacterial co-assembly versus horizontal gene transfer

Most microbial eukaryotic cultures are non-axenic and contain diverse bacterial communities. As a result, genome sequencing projects often generate data from both the target and non-target genomes. Identifying and removing contaminant contigs from these metagenome assemblies is challenging, particularly for those based only on short-read Illumina data. Illumina-only assemblies can result in many short contigs (Figure 1A) that contain one or few (sometimes fragmented) genes that may or may not belong to the target genome. In contrast, assemblies from long-read technology can produce contigs and scaffolds with hundreds or thousands of genes or even entire bacterial genomes, making it much easier to identify and remove non-target sequences from the final assembly.

Contaminant scaffolds can be identified using the Blobtools pipeline on the basis of GC content, sequencing coverage, and taxonomic assignment via BLAST searches to reference protein databases. This pipeline has been used to identify and remove contaminants from other microbial eukaryotic genome projects (Koutsovoulos *et al.* 2016; Nowell *et al.* 2018; Yubuki *et al.* 2020). During the construction of the version 2.0 assembly, we used the Blobtools pipeline to identify and remove all scaffolds from the metagenome assembly with lengths less than 1 kb or with a taxonomic assignment to bacteria, archaea, or viruses (Figure S2). These criteria resulted in the removal of 1,974 contaminant scaffolds, leaving a total of 662 scaffolds in the version 2.0 assembly (Figure 2). We also applied the Blobtools pipeline and the same filtering criteria to the version 1.0 assembly and found 99,200 contigs that were less than 1 kb in length with no taxonomic assignment and 211 contigs that were assigned to bacteria or viruses (Figure 2). Of these 211 bacterial or viral scaffolds, a majority (169, or 80%) were less than 1 kb in length, whereas 36 of them had lengths greater than 5 kb, and 21 were larger than 10 kb in length. After removing short and contaminant contigs, the size of the version 1.0 assembly was reduced to 143.4 Mb (161.8 Mb original) and 30,667 scaffolds (116,815 original) (Figure S3).

Confidently removing potential contaminant sequences has important implications for the identification of genes that arose by horizontal gene transfer (HGT) (Koutsovoulos *et al.* 2016). This is especially complicated for a group like diatoms, which are thought to contain hundreds of genes acquired by HGT from bacteria (Bowler *et al.* 2008). The version 1.0 assembly used the DarkHorse tool (Podell and Gaasterland 2007) to identify 368 foreign genes (1.7% of the 21,121 genes) from bacteria (n = 340 genes), archaea (n = 12 genes), and viruses (n = 16) (Traller *et al.* 2016). Application of our filtering routine to the version 1.0 assembly showed that 31 of the 368 HGT genes (8.4%) originally identified as foreign were located on one or more of the 211 contigs that were flagged and removed as contaminants by our filtering criteria. Repeating the Blobtools pipeline to use either 20 or 50 of the top BLASTX hits to each contig for taxonomic assignment, we flagged 540 and 699 contigs in the version 1.0 assembly as contaminants, respectively. These contaminant scaffolds contained a total 1037 and 1639 genes, respectively, with 67 (18.2%) and 73 (19.8%) of those genes present in the set of 368 HGT genes in the version 1.0 assembly.

These results show that long-read sequencing, combined with better tools to identify and remove contaminant sequences, can greatly improve genome assemblies, particularly for repeat-rich genomes that contain a mix of eukaryotic and bacterial sequences. Applying our pipeline to both assemblies, we found that the version 1.0 assembly of *C. cryptica* contained hundreds of scaffolds matching bacterial or viral proteins, whereas the version 2.0 assembly is free of contaminants (Figure 2).

**Figure 2.**
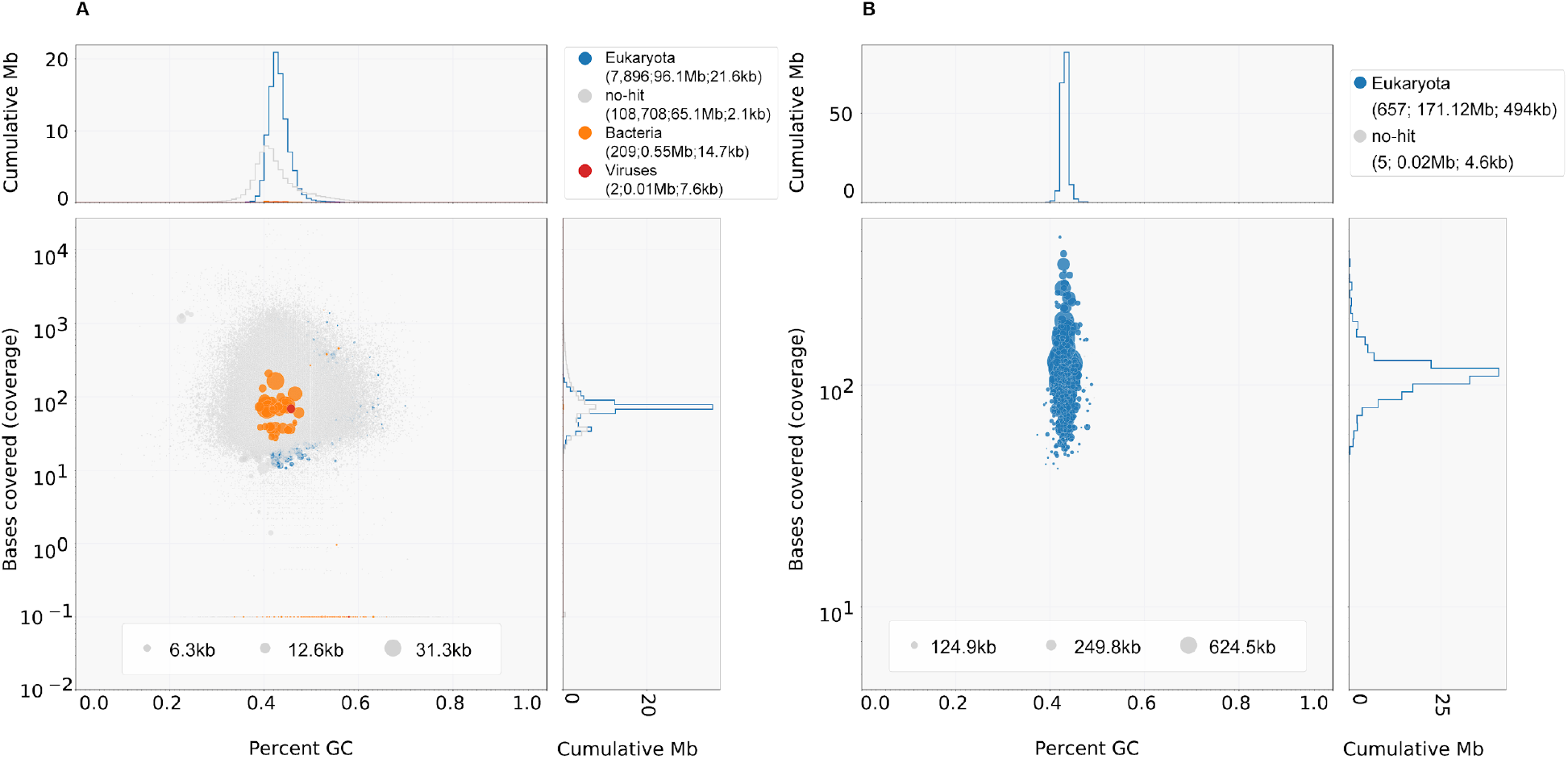
The updated assembly of *Cyclotella cryptica* is contiguous and free from contamination. Blobplots showing the taxon-annotated GC content and coverage of (A) the version 1.0 assembly, and (B) the version 2.0 genome assembly after removal of contaminant scaffolds. Legend format: “superkingdom (number of scaffolds; length of scaffolds; scaffold N50 length)”.

### Updated gene annotation of the *Cyclotella cryptica* genome

The version 2.0 assembly contains an updated and more thorough set of gene models. The updated annotation contains 21,250 gene models and 31,409 transcript isoforms (Table 2). The version 2.0 gene models contain more annotated features, including predicted genes, exons, introns, CDS (coding sequences), mRNAs (messenger RNAs), and UTRs (untranslated regions) (File S3). Our annotations of the version 2.0 assembly led to substantial increases in: (1) the mean predicted gene size [from 1.47 kb in version 1.0 to 2.09 kb in version 2.0], (2) mean exon length [608 vs. 722 bp], (3) mean intron length [125 vs. 152 bp], and (4) total length of the coding regions [27.96 vs. 41.84 Mb] (Table 2).

More importantly, we saw an increase in support for the protein gene models in the version 2.0 assembly, with a higher proportion of proteins containing Pfam protein domains (from 44.7% in version 1.0 to 46.2% in version 2.0) and matches to SwissProt (26.8% vs. 41.6%) or UniProt proteins (71.0% vs. 74.9%) (Table 2 and Table S4). These increases were possibly due to longer lengths of transcript isoforms in version 2.0 (Table 2). We also identified 188 tRNAs and 36 ncRNAs (File S4). These updated models should better enable physiological, metabolomic, and evolutionary studies of *C. cryptica*.

Fully 96.5% of the models had AED scores less than 0.5 (Table 2), indicating that the updated gene annotations were highly concordant with the input evidence (transcripts and proteins). Additionally, the 31,409 annotated isoforms included 192/255 (75.3%) of the BUSCO conserved single-copy orthologs in eukaryotes, which represents an increase from 184/255 (72.2%) in the version 1.0 assembly (Table 2). The BUSCO counts for the updated *C. cryptica* protein models are now comparable to those of the model diatoms, *C. nana* (70.2%), *F. cylindrus* (73.7%), and *P. tricornutum* (76.5%) (Table S5).

We compared the non-redundant protein sets of versions 1.0 and 2.0 using NCBI BLASTP. Protein sets were similar overall, with 19,333 (83.2%) of the version 1.0 proteins aligned to version 2.0 proteins. Of these, 4,949 (25.6%) were perfect matches (same length, 100% identity, and full length) and 6,337 (32.8%) were the same length with high similarity (> 90% identity). The remaining 8,047 (41.6%) alignments were not the same length, but 4,221 (21.8%) of these had 100% identity.

**Table 2.**
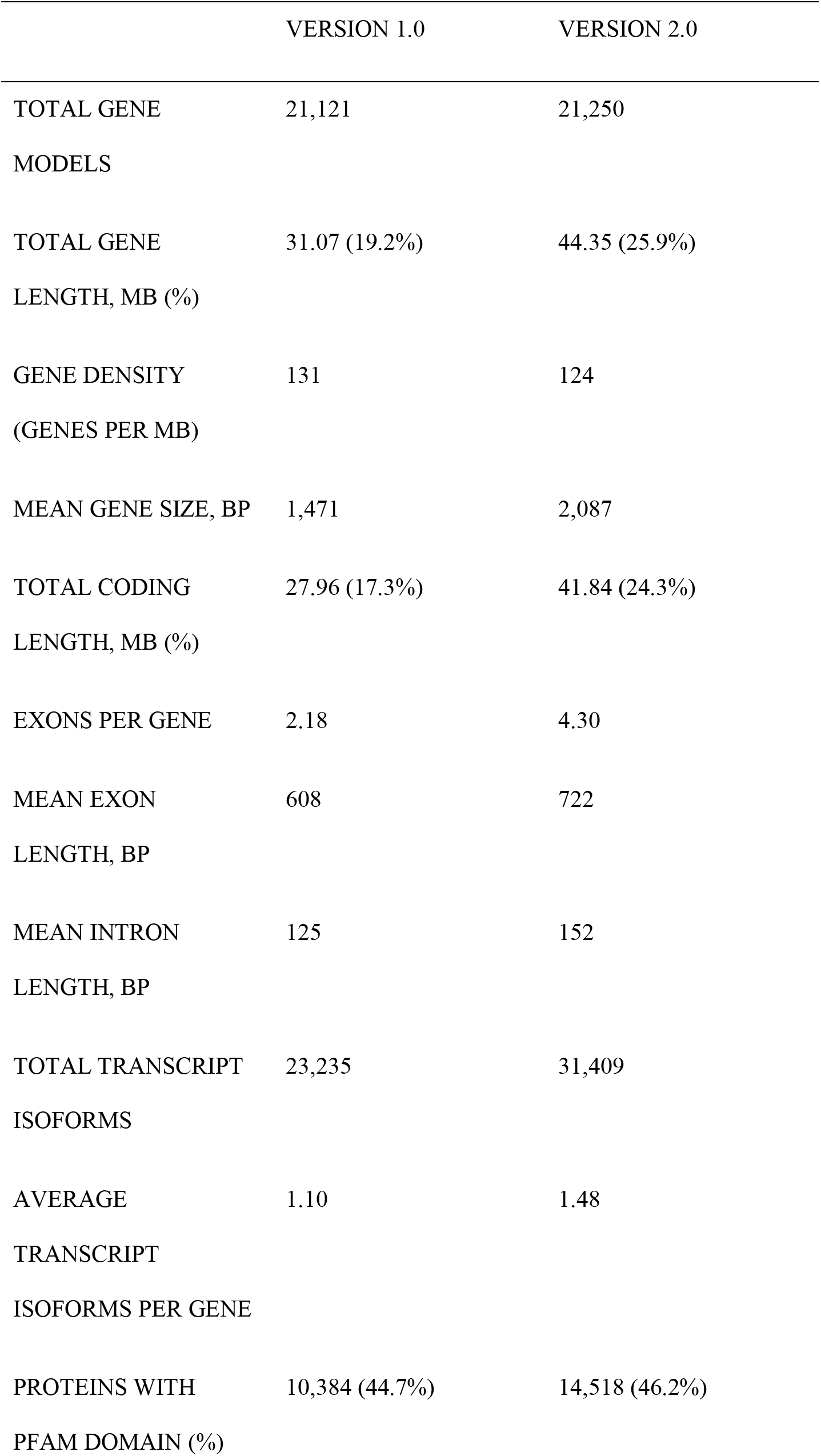
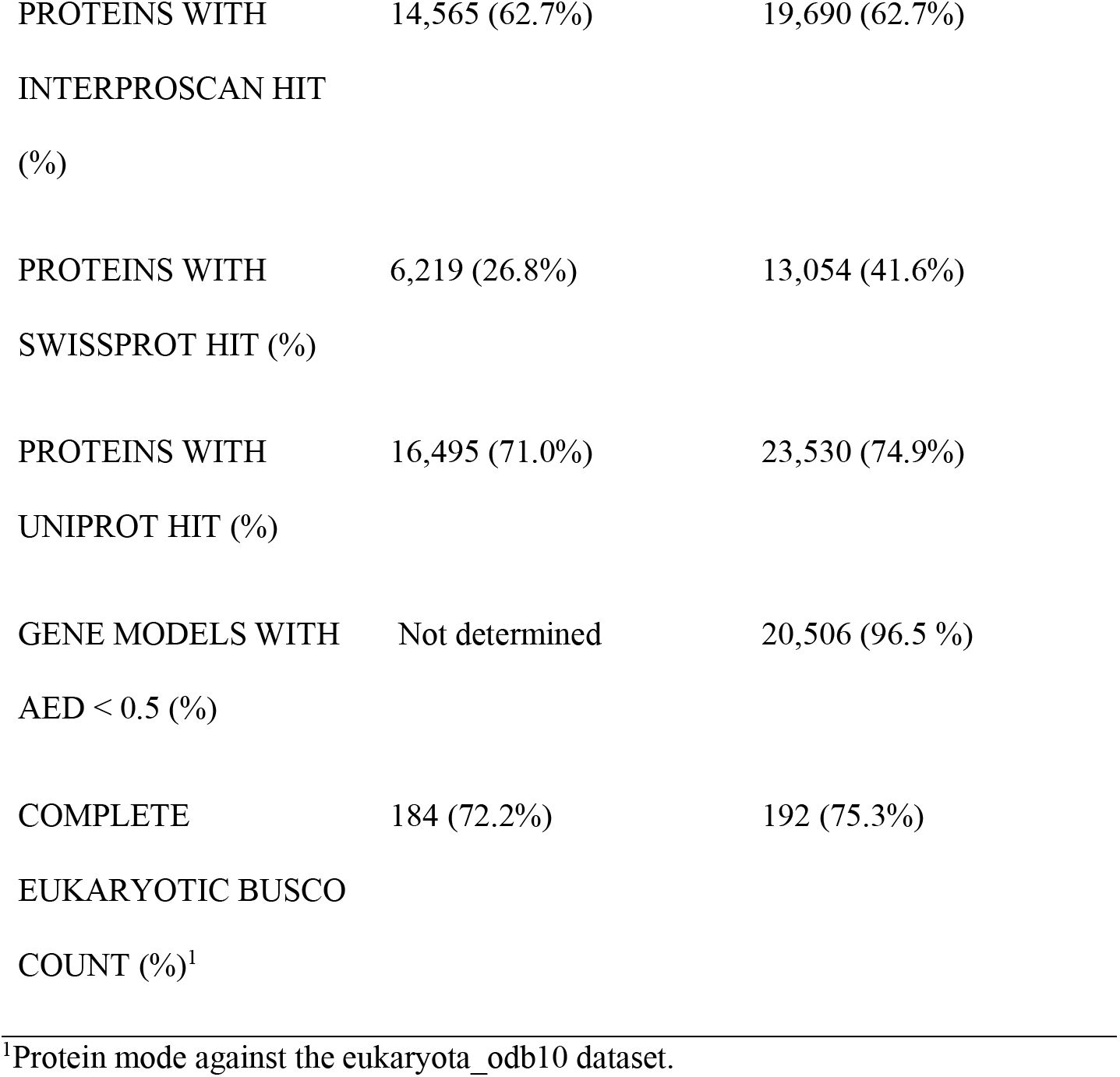
Summary of the *Cyclotella cryptica* genome annotations.

### Repeat landscape of the *Cyclotella cryptica* genome

We also revisited the characterization of repetitive elements in the *C. cryptica* genome by applying more robust structural and *de novo* discovery approaches (File S5). Repeats collectively comprised 59.3% (101.5 Mb) of the version 2.0 assembly, which was slightly greater than the version 1.0 assembly (53.8%, 98.3 Mb) (Table 1 and Figure 3). We also classified a greater fraction of the genome as transposable elements (TEs) in the version 2.0 (32.4%) than version 1.0 (12.9%) assemblies (Figure 3). Additionally, the number of unclassified repeat elements decreased from 40% to 24% between the version 1.0 and version 2.0 assemblies (Figure 3). Repeats represent just 2% and 12% of the genomes of *C. nana* and *P. tricornutum* (Armbrust *et al.* 2004; Maumus *et al.* 2009; Rastogi *et al.* 2018) (Table 1).

Among Class I retrotransposons, we identified short interspersed nuclear elements (SINEs), long interspersed nuclear elements (LINEs), and long terminal repeats (LTRs) (Figure 3 and Table S6). SINEs were not identified in the *C. cryptica* version 1.0 assembly and have only been identified in later annotations of the *P. tricornutum* genome (Rastogi *et al.* 2018). SINEs are known for their impacts on mRNA splicing, protein translation, and allelic expression (Kramerov and Vassetzky 2011) and were previously thought to be absent from unicellular eukaryotes (Kramerov and Vassetzky 2011). Their functional roles, if any, in diatoms remain poorly understood. Similar numbers of LINEs were identified in versions 1.0 and 2.0 genomes (2,626 vs. 2,350) (Table S6). We detected fewer numbers of LTRs in *C. cryptica* version 2.0 than version 1.0 (26,418 vs. 43,176), but these elements appear to represent a larger fraction of the genome (20.9%) than previously thought (8.6%) (Figure 3 and Table S6). Comparative genomics has established that diatom genomes contain diatom-specific Copia-like LTR elements called CoDis (Maumus *et al.* 2009). Gypsy-type LTR elements were predominant in *C. cryptica* and covered approximately 22.7 Mb of the genome, whereas Copia-type LTR elements covered 9.5 Mb (Table S6). Both Gypsy- and Copia-type LTRs have been identified in *C. nana* (Armbrust *et al.* 2004; Maumus *et al.* 2009), whereas only Copia-type LTRs have been found in *P. tricornutum* (Rastogi *et al.* 2018).

We also identified higher numbers of Class II DNA transposons in the version 2.0 assembly than version 1.0 (52,786 vs. 15,402), constituting a higher proportion of the genome (11.5%) than the previous assembly (3.2%) (Figure 3 and Table S6). These elements in the version 2.0 genome were classified into 12 superfamilies: *Crypton, Ginger, EnSpm, hAT, Helitron, Kolobok, MuDr, PiggyBac, PIF-Harbinger, Polintron, Sola*, and *TcMar* (Table S6). The age distribution of TEs, based on sequence divergence from exemplar elements in the repeat library, indicates that there has been a steady accumulation of DNA TEs over time in the *C. cryptica* genome (Figure 3). In comparison, DNA TEs make up less than 1% of the genome in both *C. nana* and *P. tricornutum* (Maumus *et al.* 2009).

With the improved genome assembly, we can infer that the large genome of *C. cryptica* is due to recent and gradual accumulation of repetitive elements, particularly LTR and DNA TEs (Figure 3), similar to the process commonly found in flowering plants (Piegu *et al.* 2006; Verde *et al.* 2013). TEs can impact gene function and regulation and may participate in the emergence of novel phenotypes (Kazazian 2004; Veluchamy *et al.* 2013). They have previously been investigated in diatoms for their roles in stress response and environmental adaptation (Maumus *et al.* 2009; Oliver *et al.* 2010; Norden-Krichmar *et al.* 2011). The expanded repeat classification of *C. cryptica* contributes to our growing knowledge of TE diversity in diatoms and their role in diatom genome evolution.

**Figure 3.**
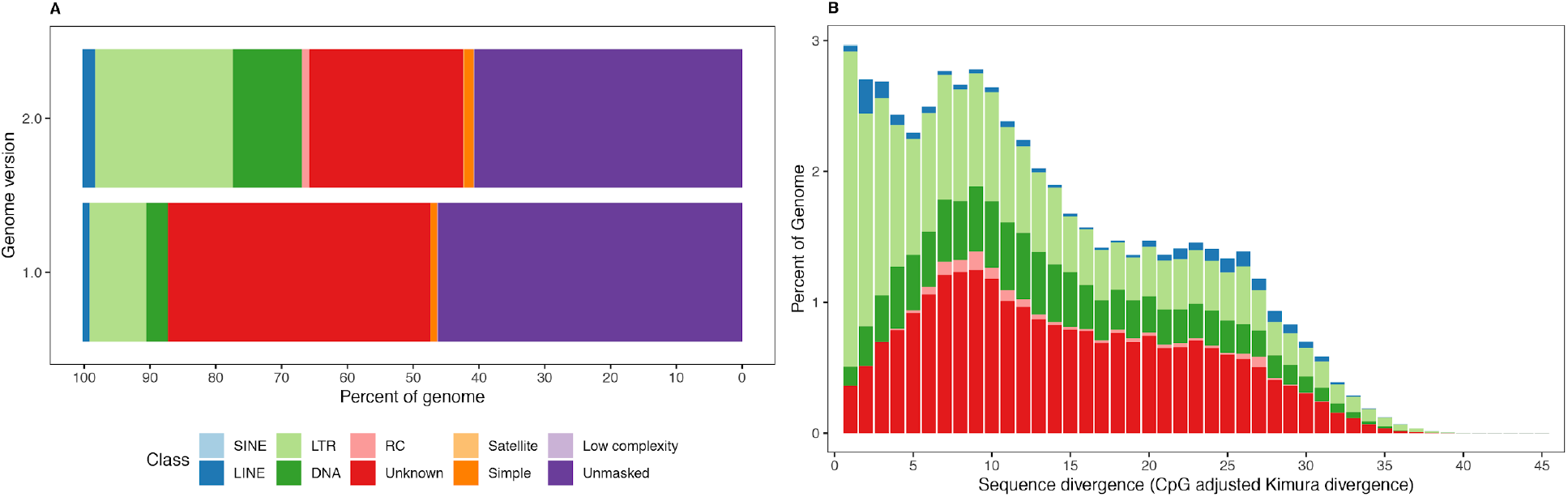
Repeat content of the *Cyclotella cryptica* genome. (A) Repeat content in the version 1.0 and version 2.0 assemblies. Bars show the proportions of the genome assemblies masked and annotated by RepeatMasker. (B) Age distribution of transposable elements in the *C. cryptica* version 2.0 genome. The total amount of DNA in each TE class was split into bins of 1% Kimura divergence, shown on the X axis (see Methods). Abbreviations: DNA, DNA transposon; LINE, long interspersed nuclear element; LTR, long terminal repeat retrotransposon; RC, rolling circle transposons (*Helitron*) SINE, small interspersed nuclear element.

## Conclusions

*Cyclotella cryptica* is one of a growing list of diatoms with a high-quality sequenced genome. The addition of long-read sequencing data improved the contiguity, completeness, and overall quality of the genome. The version 2.0 assembly allowed for new mechanistic insights into the large size of the genome, namely the historically steady and ongoing accumulation of TEs. The combination of long- and short-read sequencing data provides an effective and relatively inexpensive approach for sequencing modestly sized diatom genomes that will hopefully accelerate the pace of genomic sequencing in diatoms. The improved genome and genome annotation should also help facilitate the continued use of *C. cryptica* as a model for addressing a wide range of basic and applied research questions in diatoms.

## Acknowledgments

This work was supported by the National Science Foundation (grant no. DEB-1651087, AJA) and by a grant from the Simons Foundation (403249, AJA).

## Notes

### Competing Interest Statement

The authors have declared no competing interest.

