## Supplementary material for "Improved Reference Genome for *Cyclotella Cryptica* CCMP332, a Model for Cell Wall Morphogenesis, Salinity Adaptation, and Lipid Production in Diatoms (Bacillariophyta)": FileS2-supplemental-figures.pdf

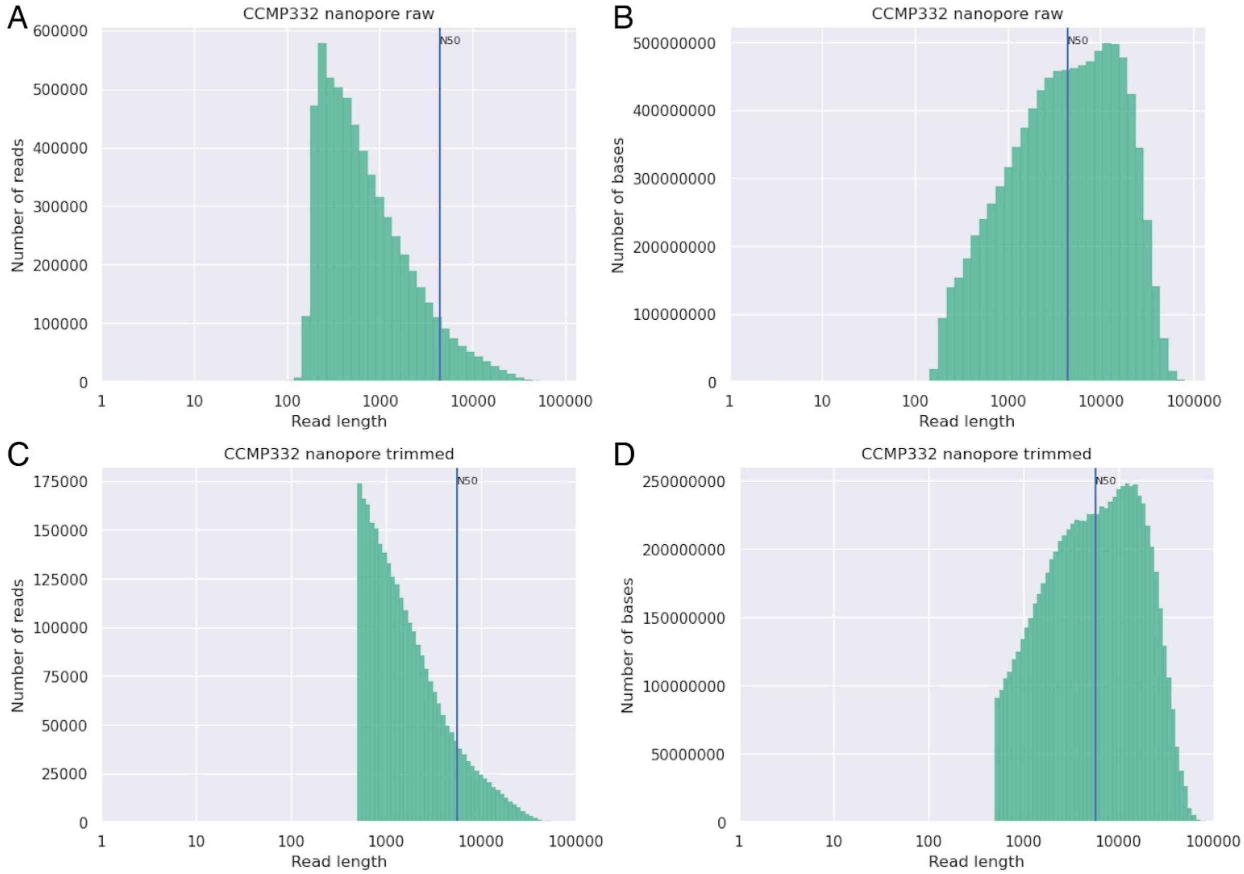


**Figure S1.** Read length histograms of the raw (A, B) and trimmed (C, D) nanopore reads.

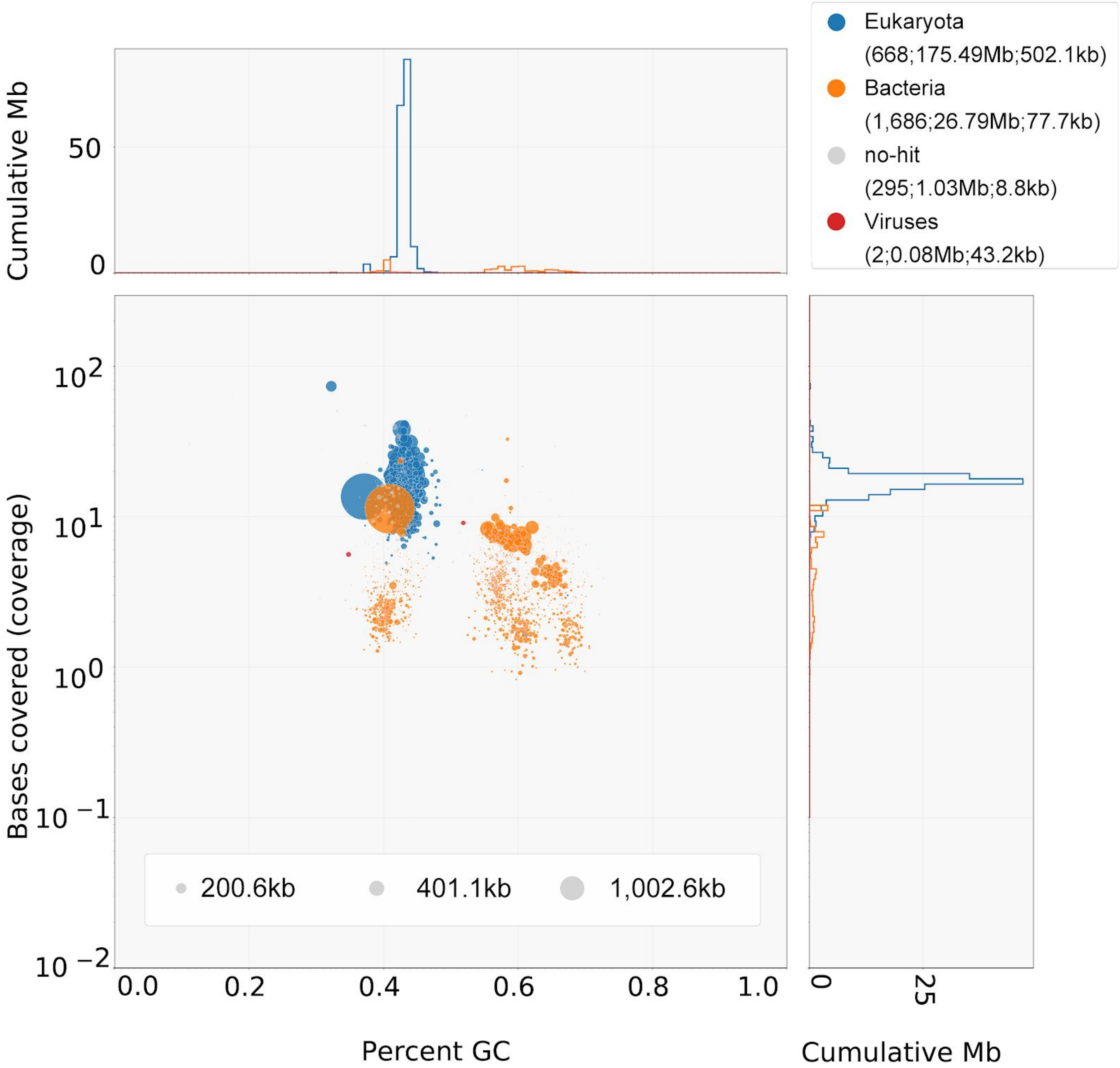


**Figure S2.** Blobplot showing the taxon-annotated GC content and coverage of the *C. cryptica* metagenome assembly. Legend format: “superkingdom (number of scaffolds; total scaffold length; scaffold N50 length)”.

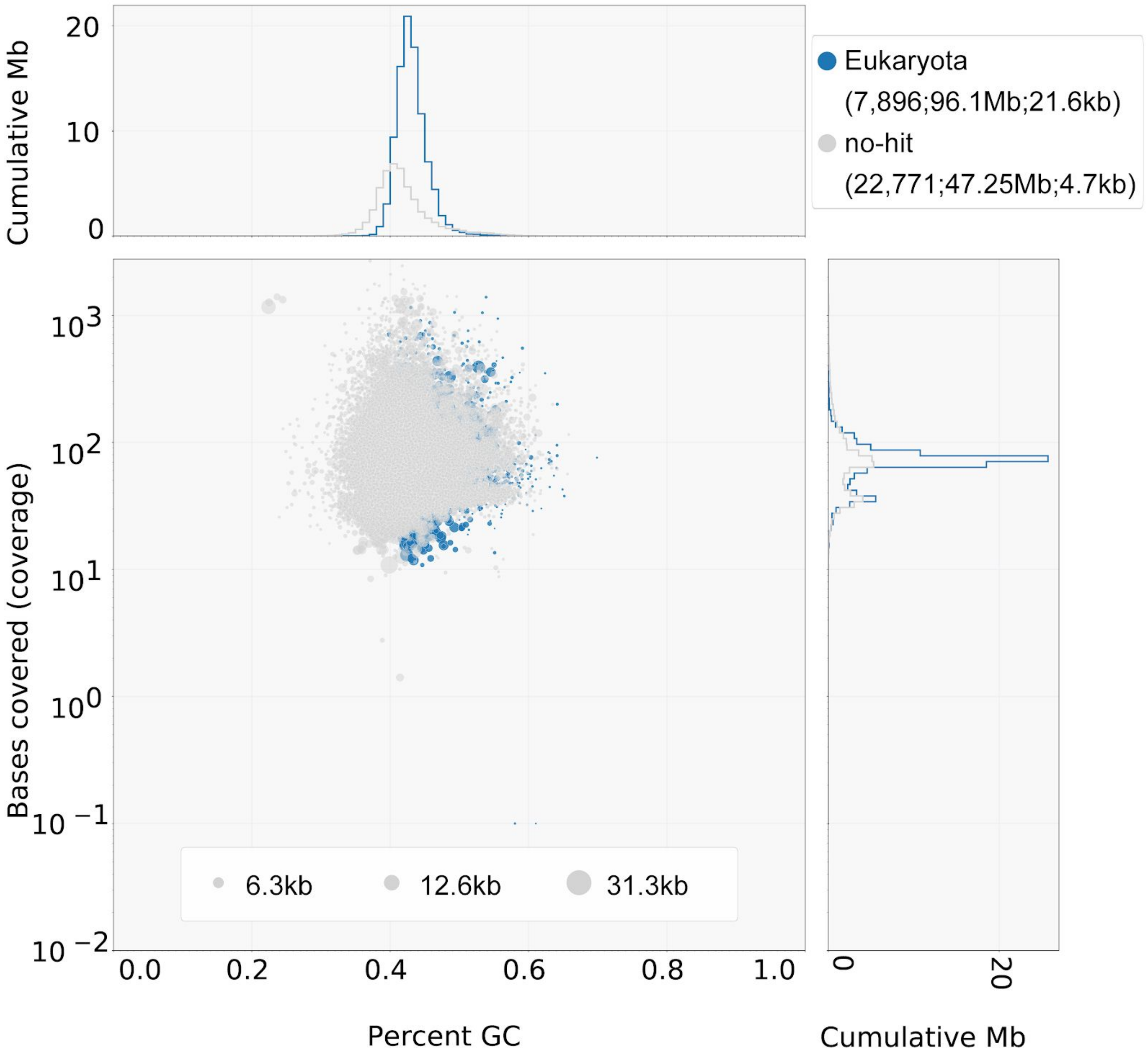


**Figure S3.** Blobplot showing the taxon-annotated GC content and coverage of the filtered *C. cryptica* ver. 1.0 assembly, after removing short (<1 kb) and contaminant scaffolds (taxonomic assignment to bacteria or viruses). Legend format: "superkingdom (number of scaffolds; total scaffold length; scaffold N50 length)".
